# The effect of above-ground vegetation on soil microbiome in urban green spaces: A systematic evidence map

**DOI:** 10.1101/2025.03.19.643203

**Authors:** Aditi Sudhir, Husnal Brar, Ananya Mukherjee, Monica Kaushik

## Abstract

Soil microbiome in urban areas is continuously shaped by urbanization and associated processes like native plant species removal, type of vegetation (native and exotic), soil physicochemical properties and anthropogenic activities, and the introduction of invasive plants, etc. Subsequently, above-ground vegetation is shaped by the soil microbiome. However, such information is hardly included while selecting plant species for greening efforts or plantations within green spaces. This could be due to a lack of studies connecting above-ground vegetation and soil microbial communities. In particular, the number of studies investigating soil microbiota and its effects on aboveground vegetation are gaining traction only recently. Existing studies vary in research questions, methodologies, urban green spaces explored, microbial community aspects, and soil characteristics examined. In this study, we are conducting a systematic evidence mapping to consolidate this research and identify global trends and gaps. We focus on the effect of above-ground vegetation on the soil microbiome in urban green spaces. Using an exhaustive search string, we retrieved 598 papers in total from two databases (Web of Science and SCOPUS) and one search engine (Google Scholar). By focusing on these relationships, we provide insights for planning and maintaining urban green spaces. This evidence mapping contributes to the scientific understanding of urban green spaces and offers practical guidance for rapidly urbanizing countries, emphasizing the integration of soil microbial communities in designing and restoring urban green spaces across different geographical locations.

## Introduction

According to the United Nations, by the year 2050, 70% of the Earth’s population will live in cities, underscoring the rapid rate of global urbanization(Sun et al., 2023). Urbanization affects biodiversity by intensifying anthropogenic activities within cities and adjoining rural and natural areas through sprawling(Filepné Kovács et al., 2024). Habitat loss and fragmentation owing to urbanization directly affect urban biodiversity, leading to lower abundance and even local extinction(Dri et al., 2021). Studies projecting urban land use expansion estimate the significant loss of terrestrial vertebrate diversity (∼30,000 species) due to habitat loss and fragmentation of natural habitats(Simkin et al., 2022; Li et al., 2022). Additionally, urbanisation indirectly affects biodiversity via increased surface and air temperature due to the urban heat island effect (Piano et al., 2019) and pollutants (Fenoglio et al., 2021). However, urban green spaces (hereafter UGS) within cities and towns provide crucial refuges for a diverse range of local biodiversity, including birds, mammals, insects, amphibians, and plants. Studies have shown that UGS supports various species, including endangered ones, by offering critical habitats amid urbanized landscapes (Soanes & Lentini, 2019). Researchers repeatedly highlight urban green spaces as essential habitats for biodiversity, supporting birds (Aronson et al., 2014), mammals (Bateman & Fleming, 2012), vegetation (Chollet et al., 2018), and insects (Baldock et al., 2015). Moreover, UGS facilitates the movement of species across fragmented urban environments, enhancing ecological connectivity and promoting genetic diversity within urban landscapes (Beninde et al., 2015). However, research on below-ground biodiversity, including microbes, is still lacking, especially in developing countries.

This may be attributed to the complex relationship that soil microbiome has with biodiversity and several other factors, many of which are elucidated in Fig.1.(see more in Fig S1). Several studies have now shown how microbial communities are interconnected with each other(Banerjee and van der Heijden, 2023). Additionally, healthy soil affects plant and animal productivity and water and air quality, thereby playing a crucial role in global air quality and water security(Banerjee and van der Heijden, 2023). Similarly, increased soil stress can eliminate organisms that aid in the discovery of new antibiotics, which is vital for a world hurtling toward antimicrobial-resistant diseases(Brevik et al., 2020). Urban soils are frequently subjected to tampering, compaction, surface sealing, and change in above-ground vegetation structure and composition, all of which will affect soil microbial diversity(Fig.1) (Fierer, 2017). Soil biogeochemistry depends on a variety of factors and varies from native systems concerning nutrient cycling, especially carbon and nitrogen(Thompson and Kao-Kniffin, 2019). Studies have shown that soil functionality depends not only on microbes but also on microbial community evenness, a lack of which affects nutrient cycling in soil, thereby affecting soil stress (Mendes et al., 2015). Additionally, a large-scale study conducted on urban green spaces and natural systems in 56 cities spanning 6 continents identified several factors, such as park management practices and environmental stress, that affect the soil microbiome(Delgado-Baquerizo et al., 2021).

**Fig. 1:**
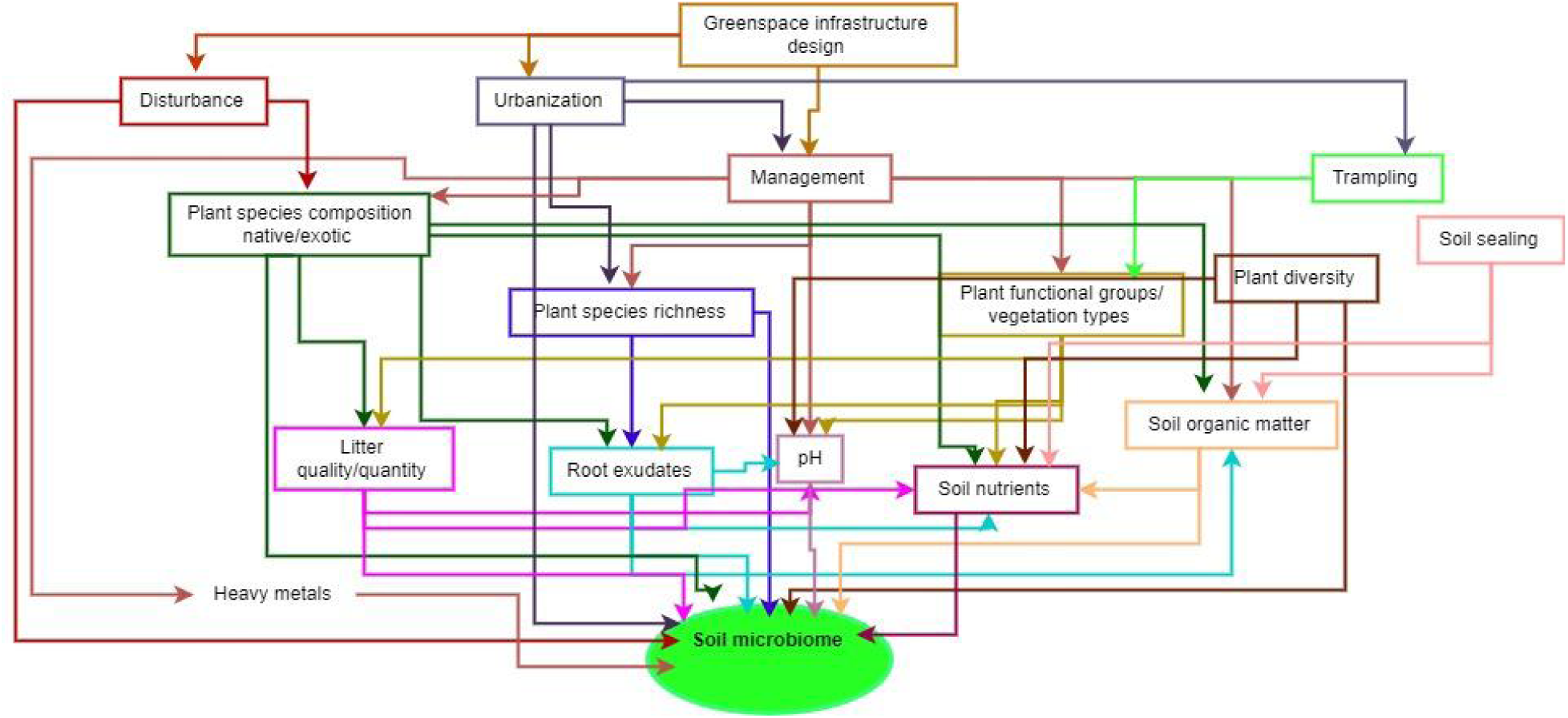
The flowchart depicts an intricate network of factors shaping the soil microbiome while highlighting the effect of above ground vegetation in the context of urban greenspaces. Most urban greenspaces face substantial disturbance and various management practices. Chosen studies have looked at different aspects of both, their effect on vegetation and hence, their effect on soil microbiome.

While in recent times, there has been an increased awareness amongst urban space managers to preserve the natural vegetation and biodiversity of such spaces, there is still a tendency to consider the aesthetic value of planted vegetation (Qiao et al., 2022). This is significant since belowground soil microbiota are rarely considered important while making such decisions, and yet they are affected by the management of land use systems(Mendes et al., 2015). It is crucial to prioritize the study of soil microbial communities as they play a fundamental role in biogeochemical cycles, support plant health, enhance soil fertility, and contribute to the overall sustainability of urban green spaces.

A study of 600 soil samples collected across Central Park, New York City, revealed a high level of below-ground microbial diversity, most of which was undescribed when compared to global microbiomes (Ramirez et al., 2014). A similar study from five urban green spaces in Adelaide, Australia, showed clear associations of soil fungal diversity with above-ground vegetation, such as areas with woody vegetation, which showed the most fungal diversity(Baruch, Liddicoat, Laws, et al., 2020). A follow-up to this study showed how plant species diversity correlates with soil bacterial diversity. Both high vegetation cover, non-native species and heterogeneity in the environment increase available nutrients, thus letting a wider array of bacteria thrive (Baruch, Liddicoat, Cando-Dumancela, et al., 2020b).

Another study also saw a distinct difference in microbial communities between urban parks and natural forests (Hui et al., 2017). A global study of soil microbes across 112 sites and 6 continents revealed that urban green spaces have distinct soil bacteria and protists from natural ecosystems(Delgado-Baquerizo et al., 2021). Aside from global homogenization, urban green spaces are also hotspots of microbial activity, especially plant pathogens and fungal parasites.

Most of these studies were conducted to help city planners and urban managers plan the vegetation of the city, especially in green spaces. Based on studies so far, the soil physico-chemical properties, vegetation, and soil microbial communities are intricately linked and affect each other greatly. However, there is limited research that focuses on the effect of above-ground vegetation on soil microbial communities in urban green spaces. This study thus aims to examine the gaps in our understanding of the role played by above-ground vegetation in shaping the below-ground microbial communities. We synthesized insights from over 600 studies to highlight the factors influencing soil health and their impact on human well-being in these environments. This mapping focuses on the type of studies that have been conducted, the variety of vegetation that is planted, and the microbes that are examined. We have taken a comprehensive approach in bringing to light the different viewpoints of the studies and tried to map them to provide a better understanding of the factors that directly and indirectly affect soil microbial diversity in urban green spaces due to above-ground vegetation.

## Objectives

The primary questions addressed in this paper are:-

What evidence exists for the role of above-ground vegetation on soil microbiomes in urban green spaces? What are the critical gaps in the available literature?

The secondary questions are:-

1. How much research has been conducted to describe the relationship between plants and the soil microbiome in urban green spaces?
2. Which geographical locations are these studies primarily limited to, and do they adequately represent the climatic and economic variations across the globe?
3. What types of research methodologies have been conducted on urban spaces to examine the relationship between vegetation and microbes?
4. What types of microbes have been identified in these studies?

This review was conducted following a systematic approach, conforming to the ROSES reporting standards to ensure comprehensive and unbiased coverage of literature (Haddaway et al., 2018). This would also entail ensuring replicability in the methodology, allowing any academic to thoroughly understand this study and use it as a foundation for future research.

## Methods

### Search for articles

#### Search sources

We used three different databases for this map: Web of Science (WOS), SCOPUS, and Google Scholar (GS). Each search string was adapted to suit each database. The search was conducted from December 2023 to January 2024. The number of records retrieved in each string is shown in Table 1.

**Table 1:** The table consists of all the search strings used for each database and the number of records from each. The highlighted strings indicate the maximum number of records from that database.

| Databases | Search Strings | Number of Records Retrieved |
| --- | --- | --- |
| Web Of Science | ((((TS=(Urban OR pocket OR remnant OR city OR urban-rural OR settlements)) AND TS=(green space OR park OR patch OR forests)) AND TS=((soil OR pedosphere) AND (microb* OR bacteria OR fungi OR prokaryote OR eukaryote))) AND TS=(plant AND (community OR diversity OR invasive OR richness OR ornamental OR exotic OR evenness OR composition) OR vegetation OR tree OR shrub) | 415 |
|  | ((((TS=(Urban OR pocket OR remnant OR city OR urban-rural OR settlements)) AND TS=(green space OR park OR patch OR forests)) AND TS=((soil) AND (microb* OR bacteria OR fungi OR prokaryote OR eukaryote))) AND TS=(plant AND (community OR diversity OR invasive OR richness OR ornamental OR exotic OR evenness OR composition) OR vegetation) | 349 |
|  | ((((TS=(Urban OR pocket OR remnant OR city OR urban-rural OR settlements)) AND TS=(green space OR park OR patch OR forests)) AND TS=((soil OR pedosphere OR turf) AND (microb* OR bacteria OR fungi OR prokaryote OR eukaryote))) AND TS=(plant AND (community OR diversity OR invasive OR richness OR ornamental OR exotic OR evenness OR composition) OR vegetation) | 417 |
|  | ((((TS=(Urban OR pocket OR remnant OR city OR urban-rural OR settlements)) AND TS=(green space OR park OR patch OR forests)) AND TS=((soil OR pedosphere OR turf) AND (microb* OR bacteria OR fungi OR prokaryote OR eukaryote OR belowground biodiversity))) AND TS=(plant AND (community OR diversity OR invasive OR richness OR ornamental OR exotic OR evenness OR composition OR native) OR vegetation OR tree OR shrub) | 422 |
| SCOPUS | TITLE-ABS-KEY((((Urban OR pocket Or remnant OR city OR urban-rural OR settlements)) AND (green space OR park OR patch OR forests)) AND ((soil OR pedosphere OR turf) AND (microb* OR bacteria OR fungi OR prokaryote OR eukaryote))) AND (plant AND (community OR diversity OR invasive OR richness OR ornamental OR evenness OR composition) OR vegetation OR tree OR shrub) | 131 |
|  | TITLE-ABS-KEY((((Urban OR pocket Or remnant OR city OR urban-rural OR settlements)) AND (green space OR park OR patch OR forests)) AND ((soil OR pedosphere OR turf) AND (microb* OR bacteria OR fungi OR prokaryote OR eukaryote OR (belowground AND biodiversity)))) AND (plant AND (community OR diversity OR invasive OR richness OR ornamental OR exotic OR evenness OR composition OR native) OR vegetation OR tree OR shrub) | 136 |
| Google Scholar | Urban OR pocket OR remnant OR city OR urban-rural OR settlements AND green space OR park OR patch OR forests AND soil OR pedosphere OR turf AND microb* OR bacteria OR fungi OR prokaryote OR eukaryote AND plant AND community OR diversity OR invasive OR richness OR ornamental OR evenness OR composition OR vegetation OR tree OR shrub | 104 |
|  | Urban OR pocket OR remnant OR city OR urban-rural OR settlements AND green space OR park OR patch OR forests AND soil OR pedosphere OR turf AND microb* OR bacteria OR fungi OR prokaryote OR eukaryote OR (belowground biodiversity) AND plant AND community OR diversity OR invasive OR richness OR ornamental OR exotic OR evenness OR composition OR native OR vegetation OR tree OR shrub | 99 |

#### Search terms and strings

We used Google Scholar to scope for articles and develop the search strings and the inclusion-exclusion criteria. The final search strings for each source used can be found below (Table 1). Web Of Science, SCOPUS, and Google Scholar were used as databases to obtain the studies for analysis. In the search string, we also used terms synonymous with cities, plants, diversity, soil, and microbiome, along with Boolean operators such as ‘AND’ and ‘OR’.

#### Supplemental searches

We identified additional articles by reviewing the bibliographies of the included studies after the primary screening. All results from each database were retrieved, and duplicate entries were removed.

#### Search limits

The scoping exercise revealed that the majority of studies began around 1990, so we included research published from 1990 to 2022. Studies from all geographic affiliations were considered, with no restrictions on location. Only studies published in English were included.

#### Estimating the comprehensiveness of the search

We estimated the comprehensiveness of the search string by checking the overlap, in terms of the number of studies, among the three databases. We found a significant overlap, indicating that all three databases covered key articles published during the selected time frame.

### Search update

These searches were conducted between December 2023 and January 2024, and no additional searches have been performed since January 2024.

#### Article Screening and Eligibility Criteria

##### Screening process

Two authors (AM, and AS) first screened the articles based on the title and abstract of the papers. The agreement coefficient between reviewers came to 0.89. The disagreement was resolved amongst reviewers to ensure a better understanding of the eligibility criteria before continuing with full-text screening. This was followed by the full-text screening. Herein, the disagreements were also resolved between the authors.

##### Eligibility criteria

Table 2 below shows the inclusion and exclusion criteria and their rationale. Although initially studies of an experimental nature were only considered, upon realizing the variety as well as the importance of other study designs, observational studies of different types and other reviews were also included.

**Table 2:**
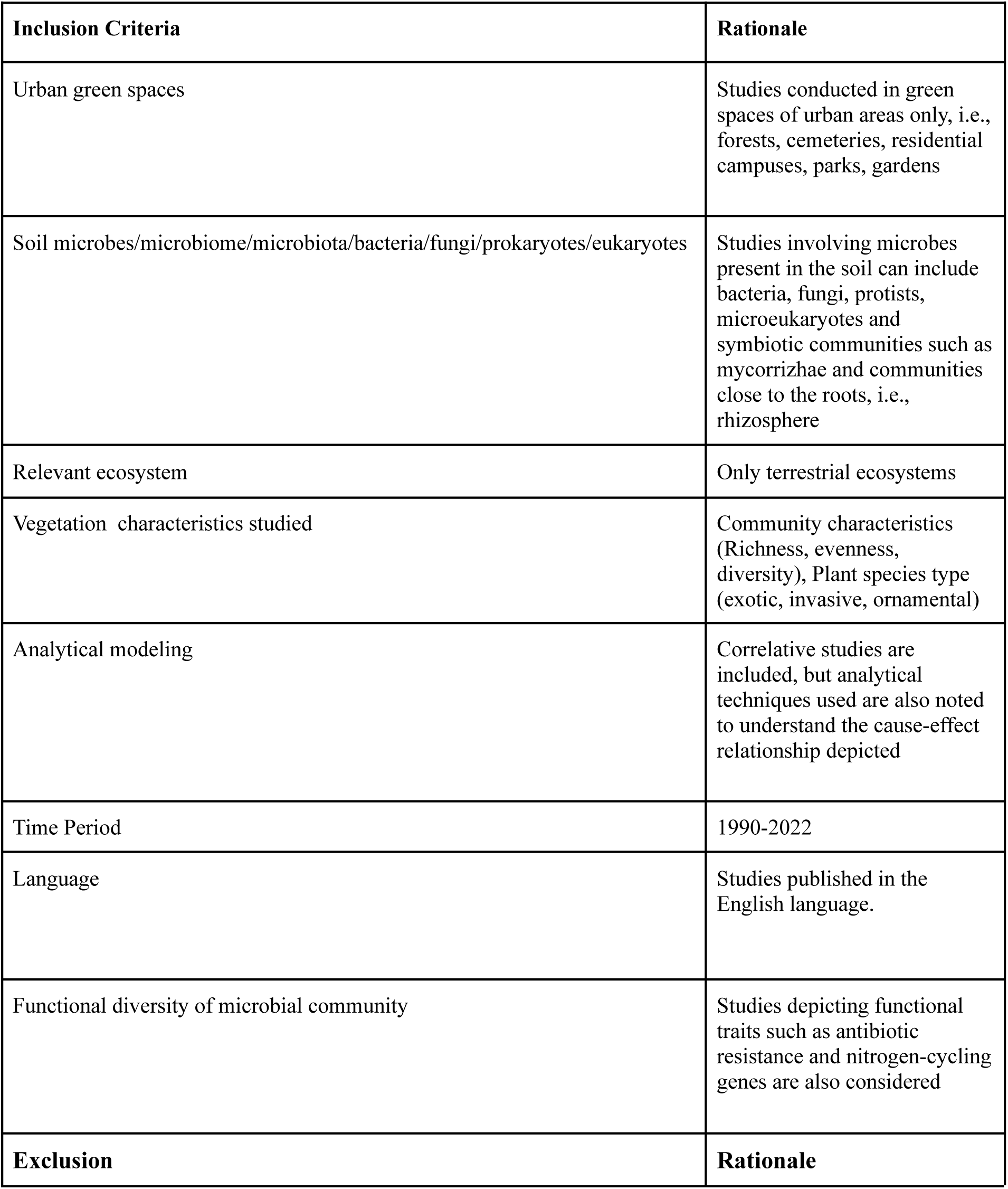
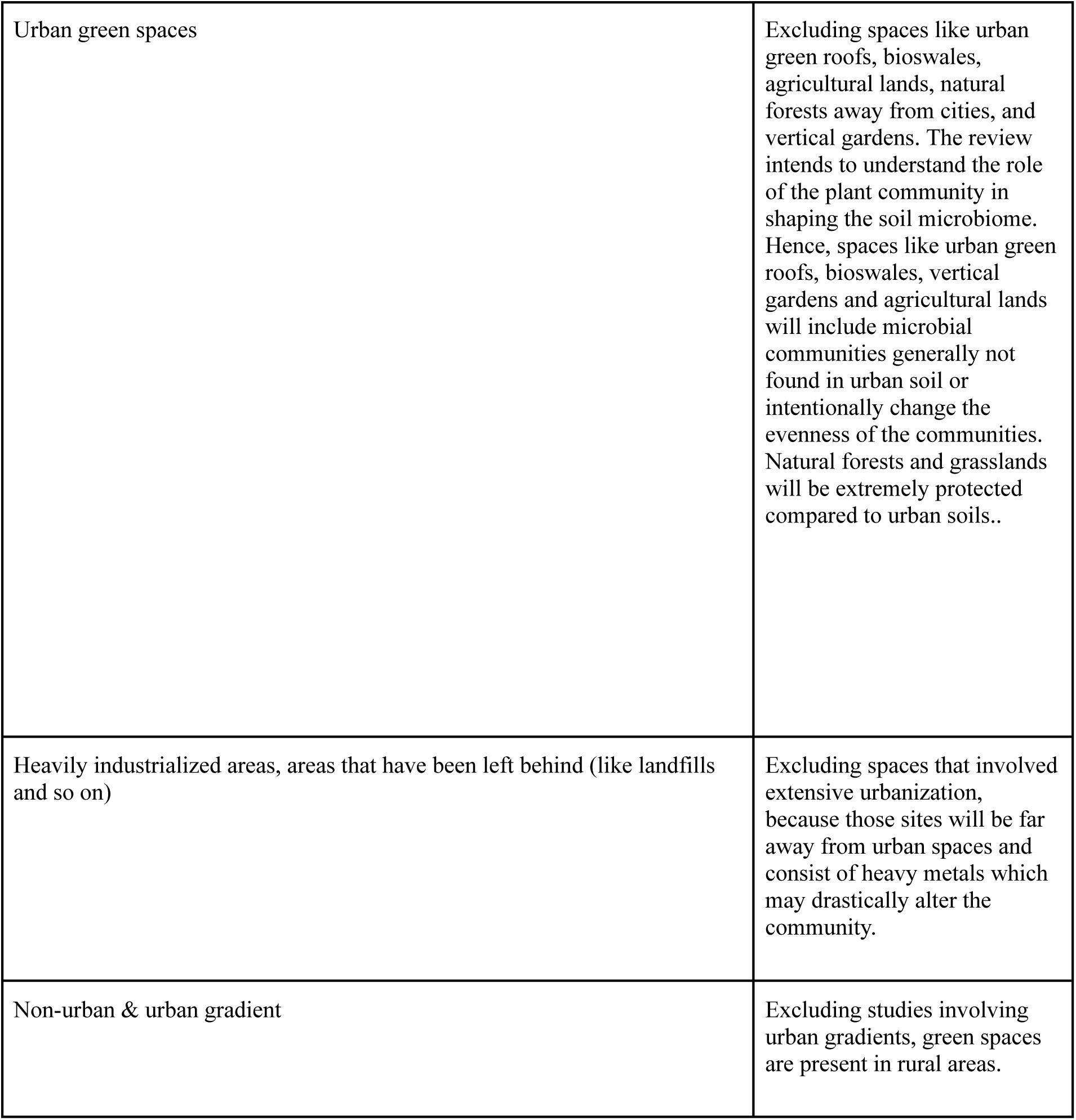
The table shows the different inclusion and exclusion criteria that were used to select the studies meant for mapping. The criteria includes synonyms and the rationale behind each criteria.

| **Inclusion Criteria** | **Rationale** |
| --- | --- |
| Urban green spaces | Studies conducted in green spaces of urban areas only, i.e., forests, cemeteries, residential campuses, parks, gardens |
| Soil microbes/microbiome/microbiota/bacteria/fungi/prokaryotes/eukaryotes | Studies involving microbes present in the soil can include bacteria, fungi, protists, microeukaryotes and symbiotic communities such as mycorrhizae and communities close to the roots, i.e., rhizosphere |
| Relevant ecosystem | Only terrestrial ecosystems |
| Vegetation characteristics studied | Community characteristics (Richness, evenness, diversity), Plant species type (exotic, invasive, ornamental) |
| Analytical modeling | Correlative studies are included, but analytical techniques used are also noted to understand the cause-effect relationship depicted |
| Time Period | 1990-2022 |
| Language | Studies published in the English language. |
| Functional diversity of microbial community | Studies depicting functional traits such as antibiotic resistance and nitrogen-cycling genes are also considered |
| **Exclusion** | **Rationale** |
| Urban green spaces | Excluding spaces like urban green roofs, bioswales, agricultural lands, natural forests away from cities, and vertical gardens. The review intends to understand the role of the plant community in shaping the soil microbiome. Hence, spaces like urban green roofs, bioswales, vertical gardens and agricultural lands will include microbial communities generally not found in urban soil or intentionally change the evenness of the communities. Natural forests and grasslands will be extremely protected compared to urban soils.. |
| Heavily industrialized areas, areas that have been left behind (like landfills and so on) | Excluding spaces that involved extensive urbanization, because those sites will be far away from urban spaces and consist of heavy metals which may drastically alter the community. |
| Non-urban & urban gradient | Excluding studies involving urban gradients, green spaces are present in rural areas. |

##### Data extraction strategy

Pilot data extraction headings were developed based on the knowledge after scoping prior to full-text extraction. As the process went on, the headings were modified to include headings that seemed relevant to our study. Any point of uncertainty throughout the process was settled after consulting with other authors. Every reasonable attempt was made to access the full-text articles for screening. The data extracted were: Geographical distribution, Study design, Vegetation identified, Microbes identified, Microbial C/N or Biomass, Type of urban green space, and the Publication date.

##### Data Mapping method

The results are presented graphically using R to develop bar graphs and a geographical map. As mentioned above, for the map metadata i.e, in vegetation identified, study design, and so on was extracted and displayed as bar graphs. The location of the studies was displayed on a geographical map. For each data point, the number of categories was decided based on the variation observed for each point. Each category has subcategories to account for the alternatives observed. The categories of each data point are as follows:

- Study design:

- Experimental
- Observational
- Review
- Systematic analysis
- Survey
- Vegetation identified:

- Composition
- Diversity
- Functional groups
- Species number
- Microbes identified:

- Prokaryotes or eukaryotes
- Based on phyla
- Microbial C/N or biomass observed:

- Microbial C and N biomass
- Bacterial biomass
- Fungal biomass
- Types of urban green spaces studies:

- Urban forests
- Urban parks
- Remnant forests and so on
- Publication date:

- Published month and year

## Results

### Number and types of articles

The search strings yielded 3,876 hits from three different databases. After the removal of duplicates, 656 articles were left, of which 536 articles were excluded after title and abstract screening. 118 articles were screened for full text, of which 10 could not be retrieved. Thus, 62 articles were included in this map. Refer to Fig.2 for details of the screening process.

**Fig. 2:**
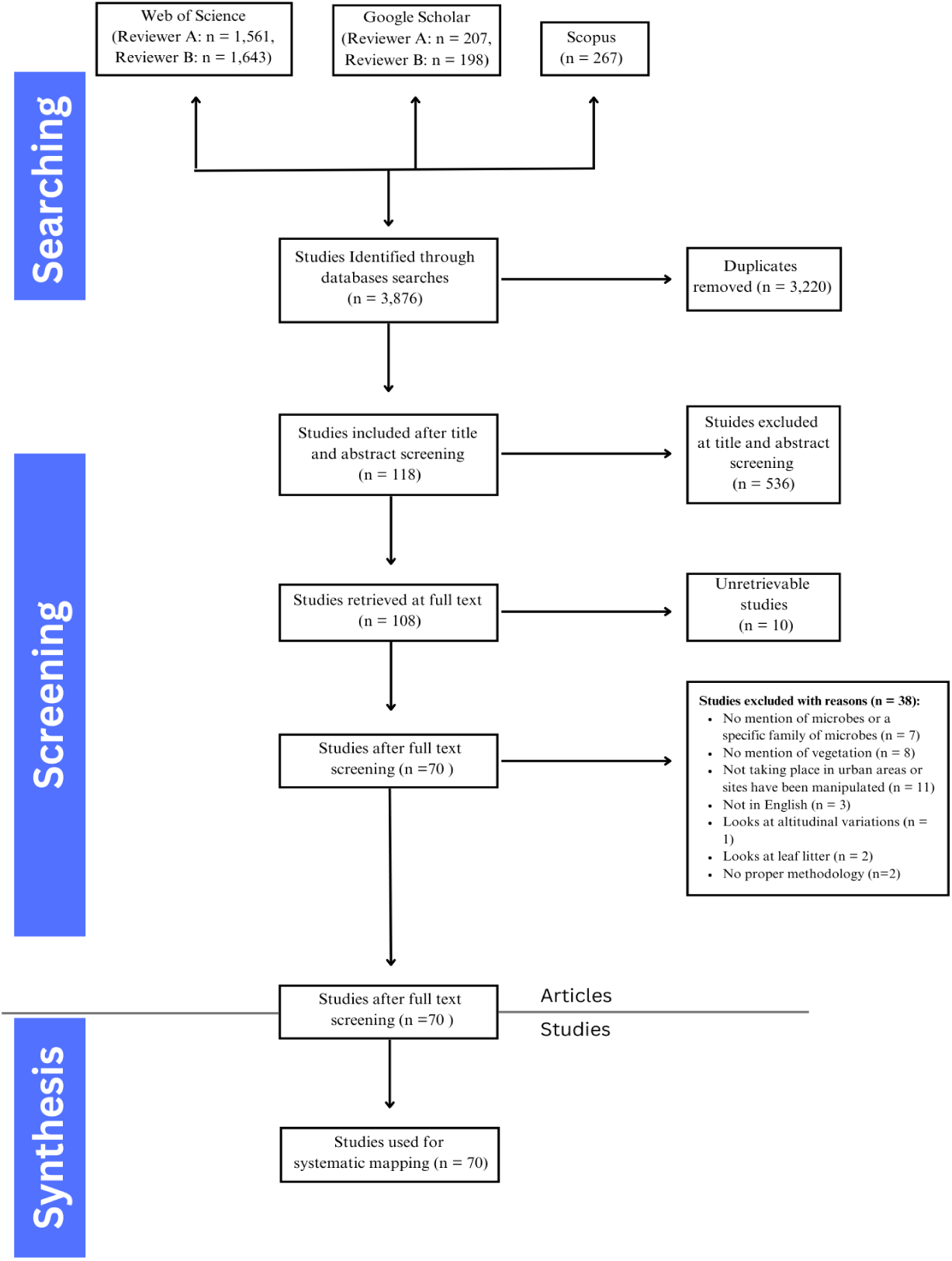
ROSES flow diagram from systematic maps (fig 3 to fig 9).The diagram shows the systematic approach taken to select the studies for mapping. It details the criteria used for selection at each step.

### Geography of included studies

The geographical distribution of the studies highlights the extent and focus of research conducted across different regions (Fig.3). The number of studies ranges from 1 to 20. The articles were not evenly divided across countries. The highest number of studies were conducted in China (n= 20), The United States of America (n= 8) and Australia (n= 6). Studies from only three developing countries were included in the evidence map. While the highest number of studies were from China, only a single study from the other two developing countries met the criteria. All other selected studies were performed in developed countries such as the USA, Germany and so on.

**Fig. 3:**
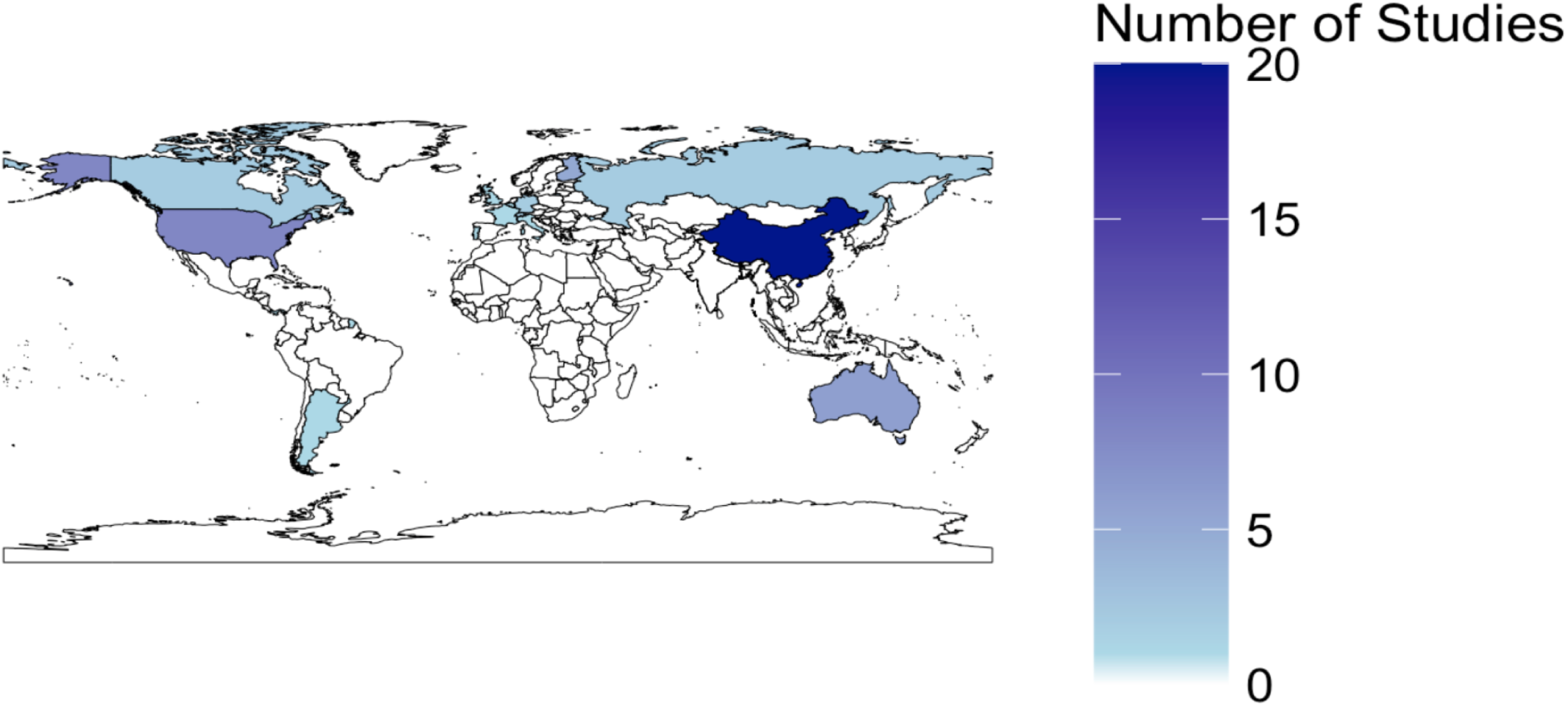
The figure maps out the geographic distribution of the shortlisted studies. Based on this, countries are highlighted according to the number of studies conducted there. As can be seen in the figure, the highest number of studies were conducted in China while very few studies are being conducted in other parts of Asia, South America and Africa.

### Type of Studies

The number and types of studies associated with the 62 publications can be seen in Fig.4. The most common type of study is the experimental study using PCR (n=34) to identify the soil microbial communities, and the least common study type was the experimental eco plate study (n=1).

**Fig. 4:**
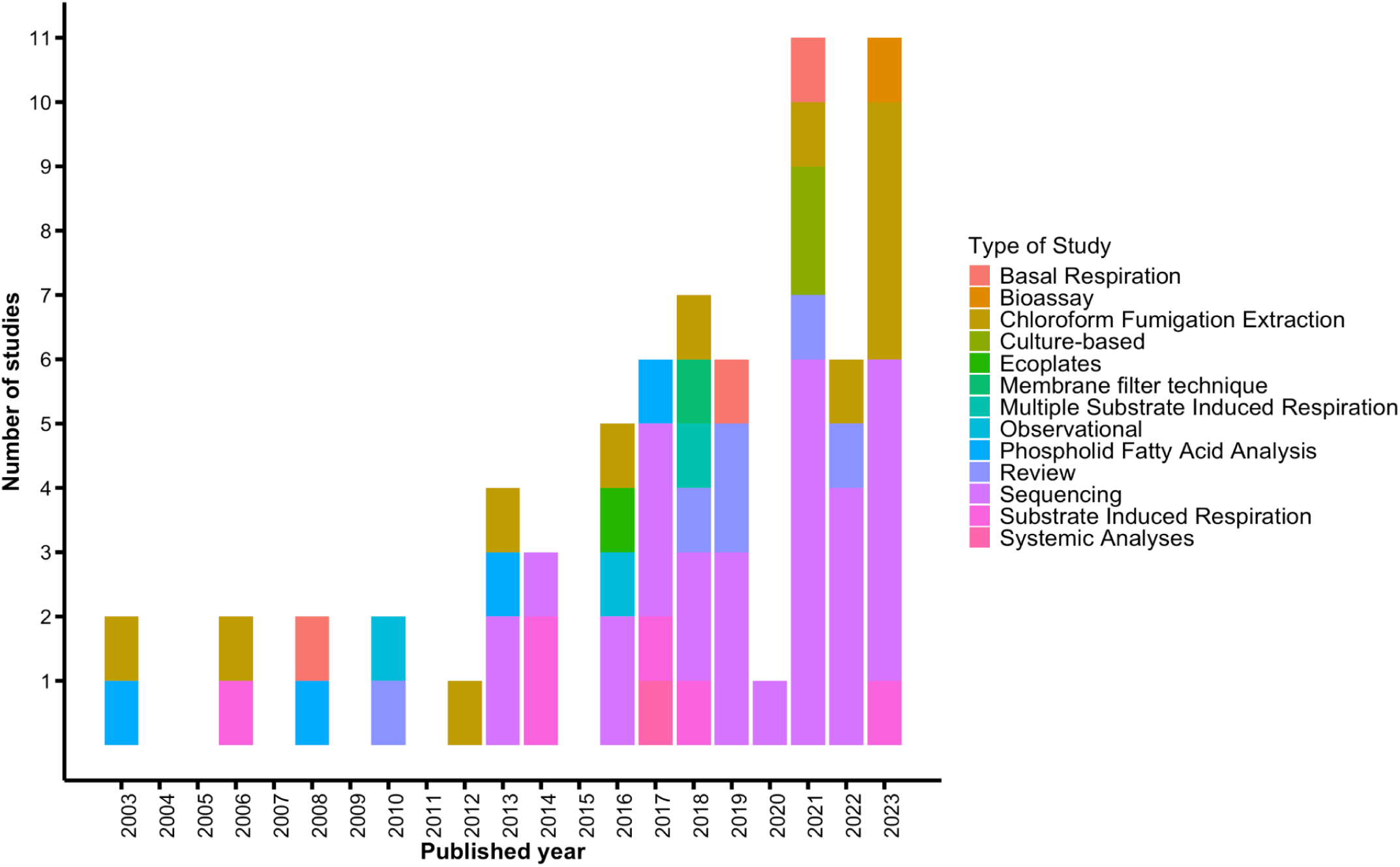
The bar chart shows the different types of studies conducted from the years 2003 to 2023. Most studies have been conducted after 2016. Maximum number of studies have used sequencing to identify microbial communities, with maximum being in 2023 and least being in 2014.

### Vegetation attributes studies

The different types of vegetation studied can be seen in Fig. 5. While the number of studies examining plant species richness, diversity and functional groups does not show a substantial difference, the number of studies considering plant species composition is significantly higher.

**Fig. 5:**
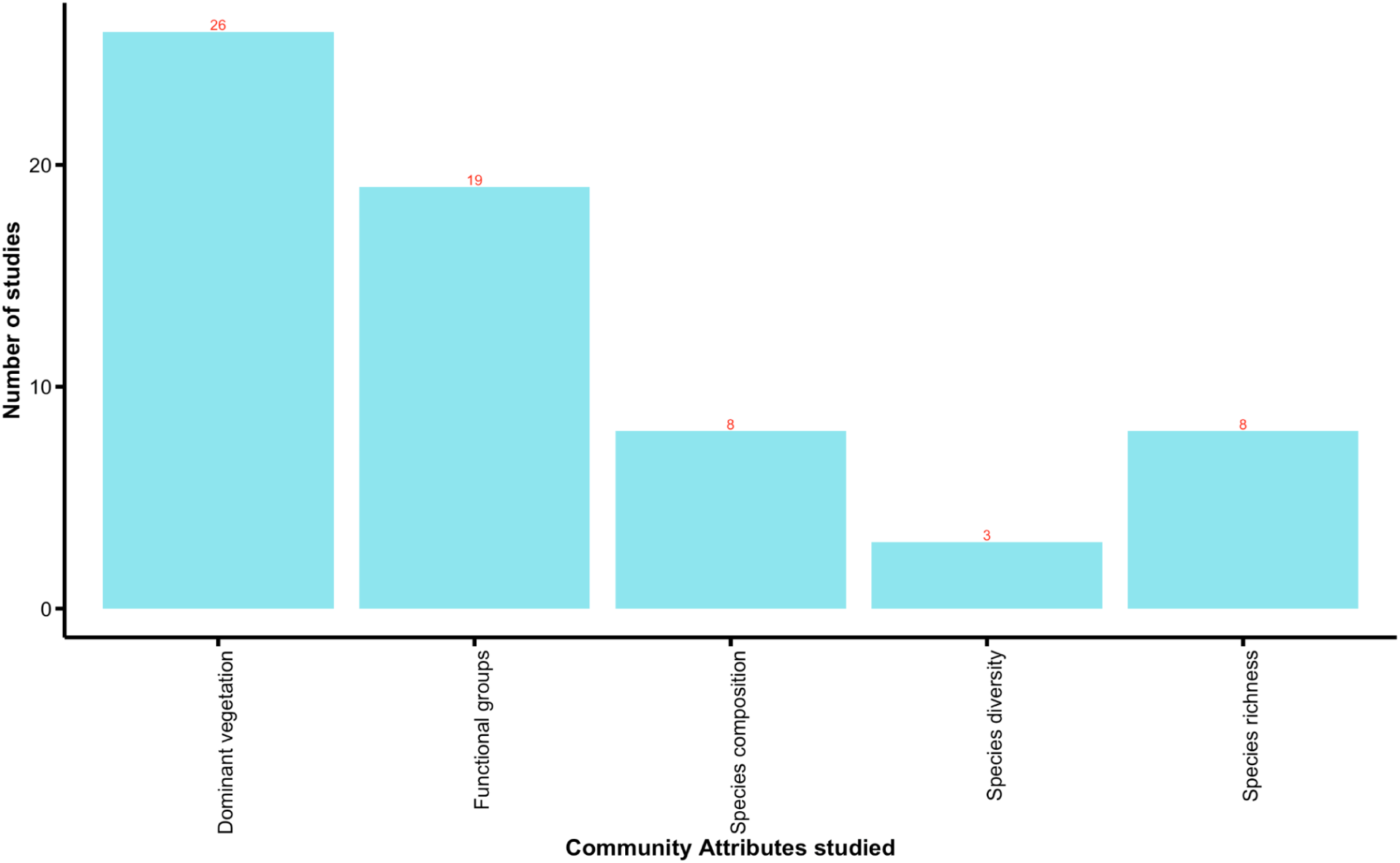
Bar chart depicts the number of studies employing different vegetation attributes to investigate the effect of vegetation on the soil microbiome.

### The type of microbes identified

Fig 6 shows the different prokaryotic genera observed. While most studies have simply mentioned the presence of bacteria. Many studies have been done on genera-level identification. Proteobacteria are the most commonly found genera, followed by Acidobacteria. While, genera such as Desulfobacterota, Elusimicrobia and so on were found the least. There were also genera belonging to Archaebacteria as well as genera such as Myxococcota which were mentioned in 6 or lesser studies, that figure is present in Supplementary Figure 1.

**Fig. 6:**
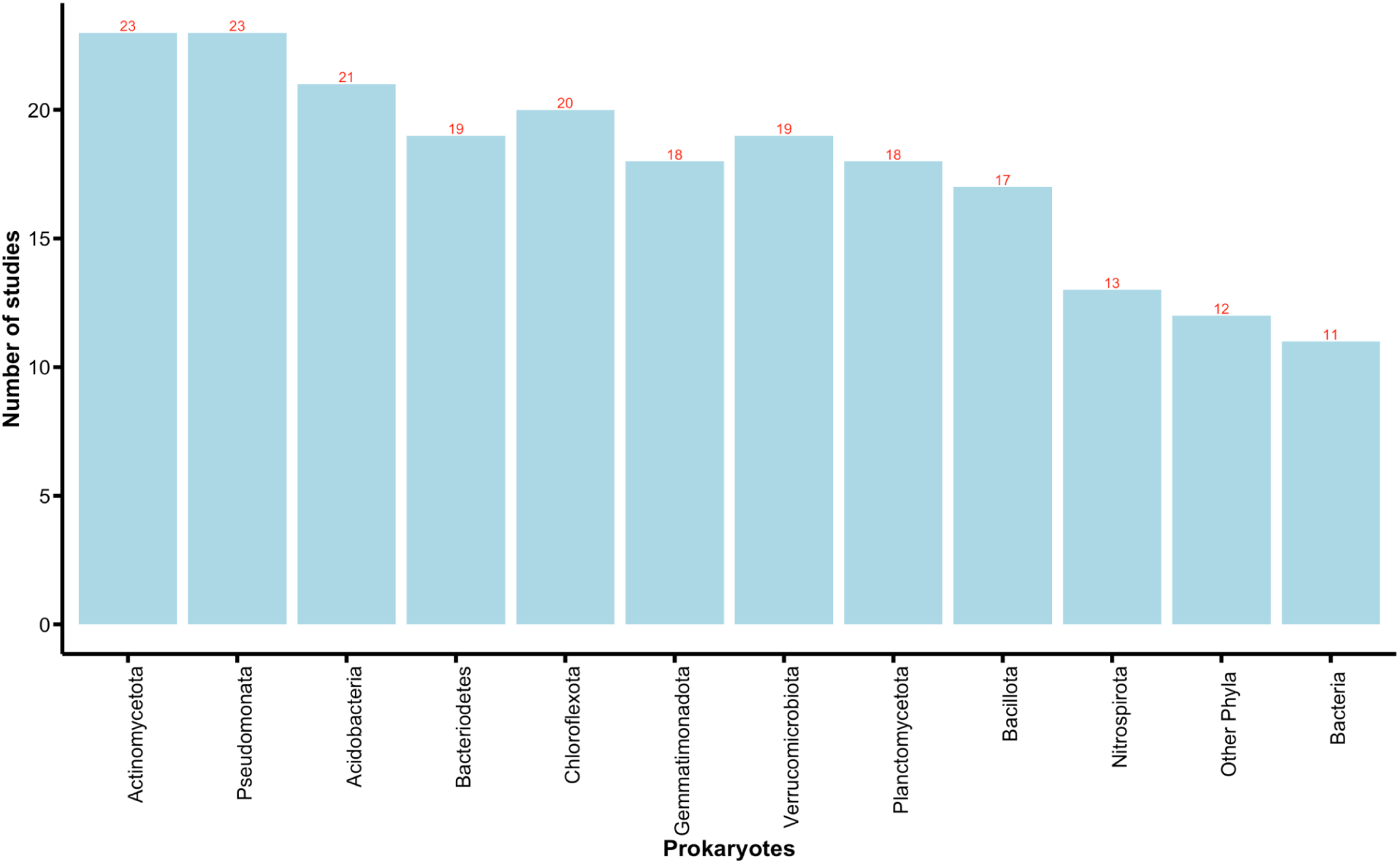
The bar chart shows the different families of prokaryotes observed across studies. The category ‘Bacteria’ are for studies that simply mention the presence of microbes belonging to these kingdoms. There were quite a few kingdoms mentioned in only 6 or lesser studies, those have been included in the supplementary.

Fig 7 shows the different eukaryotic genera observed. A considerable number of studies have simply indicated the presence of fungi. Many studies have done a genera-level identification, similar to prokaryotes. Ascomycota and Basidiomycota were most commonly found. While, genera such as Rotifera were found the least. Some studies were even able to identify some protists, i.e. microeukaryotes.

**Fig. 7:**
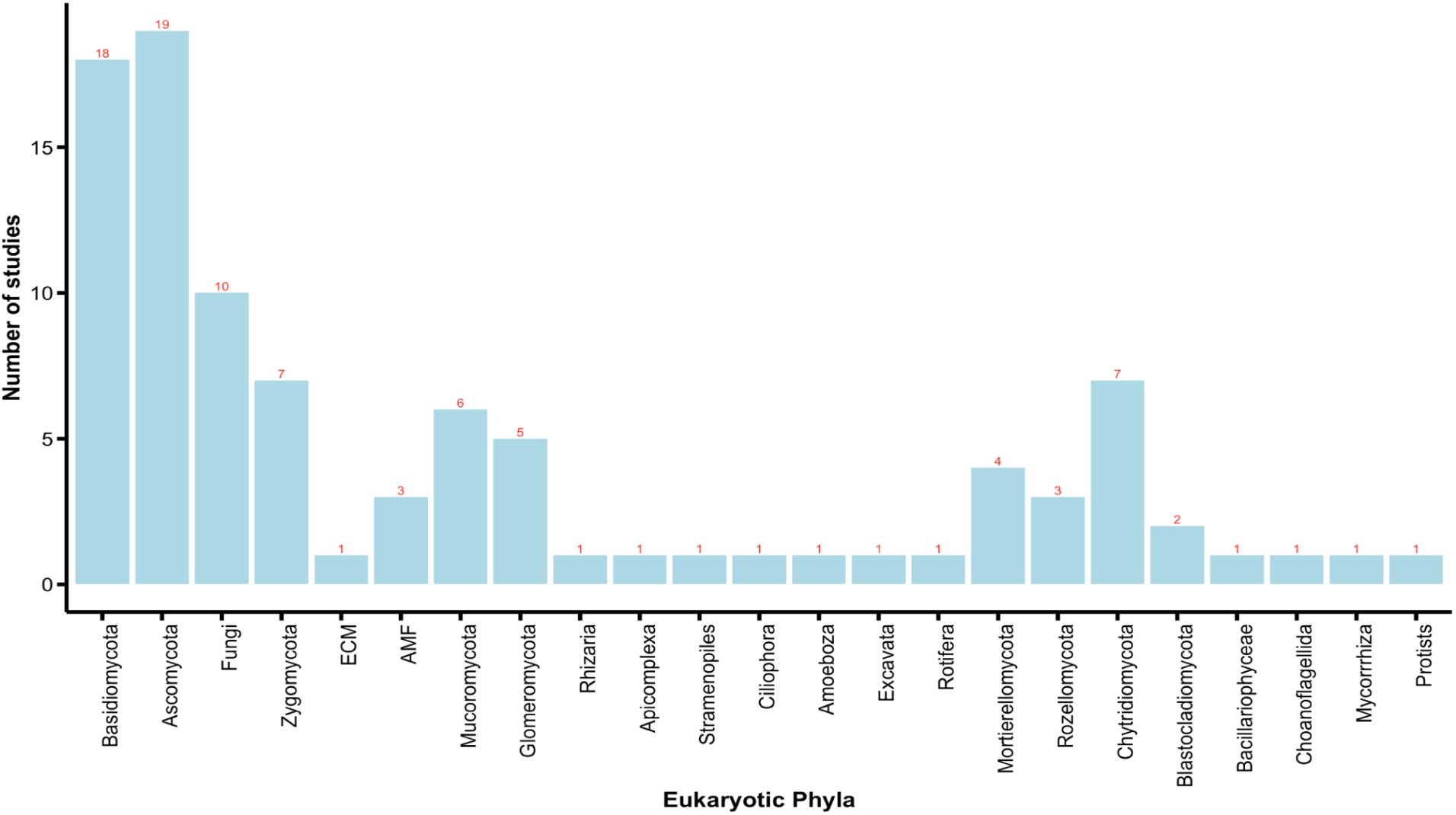
The bar chart shows the different families of eukaryotes observed across studies. The category ‘Fungi’ are for studies that simply mention the presence of microbes belonging to these kingdoms.

### Studies that show microbial biomass

Fig. 8 shows the number of studies across publications that show microbial biomass. Whilst most studies resort to sequencing (as can be seen in Fig. 4), a few choose to study biomass. From those, most studies have chosen to observe microbial, carbon, and nitrogen biomass. However, very few (n=4) have used it for further identification.

**Fig. 8:**
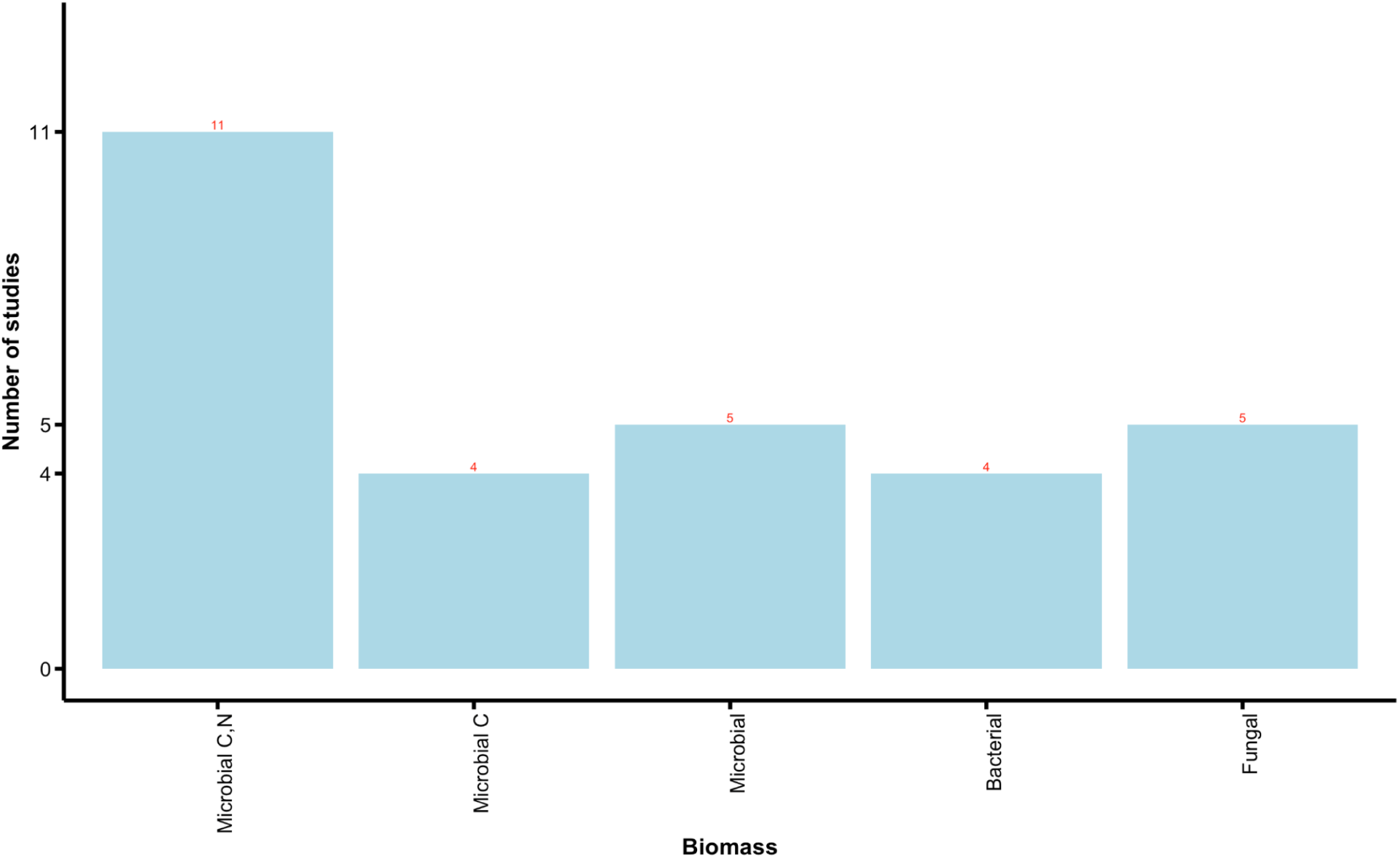
The bar chart shows the number of studies that chose to study and identify through biomass, instead of sequencing.

### Types of urban green spaces

Fig. 9 shows the different types of urban green spaces the studies are spread across. Most studies (n=36) have observed microbial diversity in urban parks and forests. While, areas like vacant lots, home gardens, and private backyards have been studied the least.

**Fig. 9:**
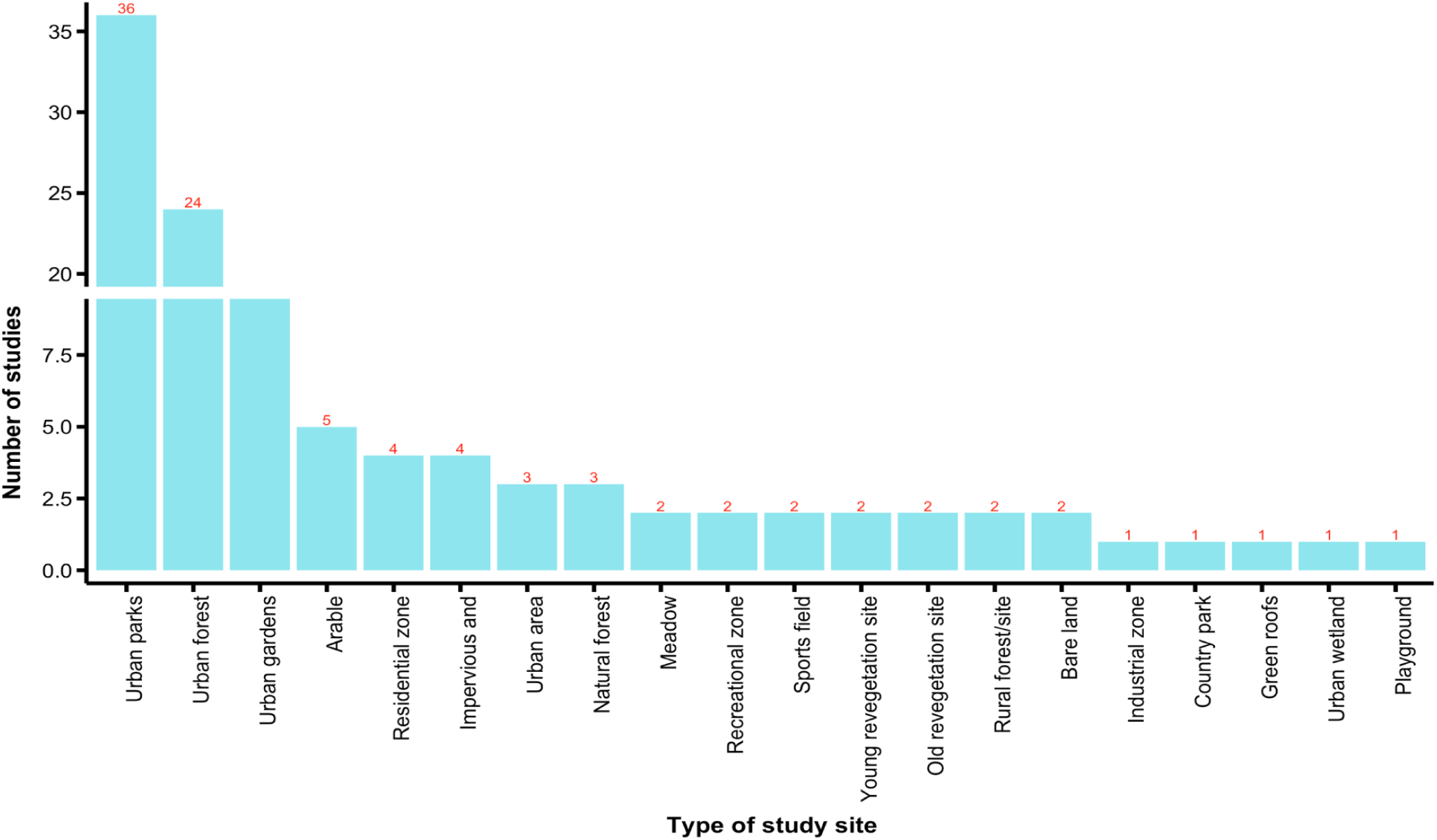
The bar chart shows different urban greenspaces studied across the included studies.

## Discussion

This study offers a comprehensive synthesis of current knowledge on the influence of above-ground vegetation on below-ground soil microbial communities in urban green spaces. By systematically mapping the existing research, we identify critical gaps and biases that hinder a holistic understanding of these interactions. Notably, there is a glaring lack of research from biodiversity-rich regions such as Africa, India, and Southeast Asia (Fig. 3), limiting our ability to generalize findings globally. Additionally, existing studies have disproportionately focused on natural forest fragments to explore the effects of above-ground vegetation on soil microbial communities, while other urban green spaces, such as parks and playgrounds, remain underexplored (Fig. 8). Addressing these disparities is essential to develop a more inclusive and comprehensive framework for understanding soil-plant-microbe interactions in urban ecosystems.

The geographical distribution of studies on the soil microbiome reveals some noteworthy patterns (Fig.3). A significant concentration of research originates from developed countries such as the United States, Germany, and other European nations, reflecting their advanced research infrastructure and long-standing focus on ecological and environmental sciences (Delgado-Baquerizo et al., 2018; Vasar et al., 2022). Interestingly, China is a leading contributor, with the highest number of studies in this field. This trend suggests a growing recognition in China of the soil microbiome’s importance, likely driven by its integration into national policies on conservation, agriculture, and public health (Y. Wang et al., 2024; Yongguan et al., 2017). Despite these advancements, there remains a critical gap in research from biodiversity-rich yet underrepresented regions such as parts of Africa, South America, and Southeast Asia, where restoration efforts and conservation initiatives are urgently needed (Guerra et al., 2020; Díaz et al., 2021). Given the pivotal role that soil microbiomes play in shaping above-ground vegetation and ecosystem resilience, addressing this geographical imbalance is crucial. Expanding research efforts into underrepresented regions will not only provide valuable insights into the microbial diversity of these areas but also inform sustainable management and restoration strategies in the face of global challenges such as deforestation, climate change, and biodiversity loss (Peddle et al., 2024; Bertola et al., 2021b).

Figure 4 highlights the evolution of methodologies used to study the soil microbiome from 2003 to 2023. Early studies (2003–2010) relied on techniques like Chloroform Fumigation Extraction (CFE) and Substrate Induced Respiration (SIR) to estimate microbial biomass, offering limited insights into diversity. Around 2012, the adoption of sequencing methods began rising, becoming dominant by 2015. This aligns with studies such as Fierer and Jackson (2006b) and Lauber et al. (2009b), which demonstrated sequencing’s ability to reveal microbial diversity and biogeographical patterns. While culture-based methods persist, their use has diminished due to known biases (Nannipieri et al., 2003). Post-2015, methods like PLFA analysis and systemic analyses gained prominence, reflecting a focus on linking microbial communities to ecosystem functions (Six et al., 2006; Delgado-Baquerizo et al., 2016b). The peak in methodological diversity around 2020–2021 indicates the rise of interdisciplinary approaches, including multi-omics, which have advanced our understanding of microbial roles in ecosystem processes (Jansson & Hofmockel, 2019).

The most studied community attribute was dominant vegetation (26 studies), while species diversity was the least studied (3 studies) (Fig. 5). Studies on dominant vegetation provide a baseline understanding of ecosystems but often neglect the effects of diverse plant communities, which rarely exist as monocultures (Wardle et al., 2004). Attributes like functional groups (19 studies) and species richness (8 studies) highlight the growing focus on plant traits and biodiversity, known to shape belowground processes (Tilman et al., 1997; Loreau et al., 2001). Underexplored attributes such as species diversity and composition (11 studies combined) are critical for understanding microbial community structure and resilience, as species-rich ecosystems promote greater microbial diversity and functional stability (Hooper et al., 2005; (Van Der Heijden et al., 2007).

Most studies predominantly reported or commonly observed major prokaryotic phyla, including Actinomycetota and Pseudomonadota (Fig 6). These two groups play crucial ecological roles in soil formation. Actinomycetota are known for their ability to degrade complex organic compounds, contributing significantly to decomposition processes and nutrient cycling in soils, particularly in managed or urban soils (Barka et al., 2016)). Pseudomonadota, on the other hand, are versatile and often function as plant-growth promoters and biocontrol agents, with their adaptability allowing them to thrive in diverse soil environments (Lugtenberg & Kamilova, 2009). While these major phyla are more adaptable and generalist in nature, the minor phyla reported may reflect specific microbial communities associated with unique vegetation at the study sites. These minor groups might not be native to the soil but could be transient populations influenced by local vegetation and environmental factors (Fierer & Jackson, 2006: Delgado-Baquerizo et al., 2018).

Among eukaryotes, fungal phyla dominated the studies (Fig. 5), with Ascomycota (19 studies) and Basidiomycota (18 studies) receiving the most attention. These groups play key roles in nutrient cycling, decomposition, and symbiotic relationships with plants (Peay et al., 2016). Other fungal groups, such as Zygomycota, Chytridiomycota, and Mortierellomycota, were less frequently studied, likely due to their narrower ecological roles or detection challenges (Blackwell, 2011). Moderate attention to Glomeromycota (5 studies) and arbuscular mycorrhizal fungi (AMF) (6 studies) reflects their significance in plant nutrient uptake and stress tolerance (Van Der Heijden et al., 2015). Non-fungal eukaryotic groups, including Rotifera (4 studies) and Rozellomycota (3 studies), were less frequently examined, and phyla like Apicomplexa and Stramenopiles appeared in only a single study each. The dominance of fungal studies highlights their central role in plant-microbe interactions, but the underrepresentation of non-fungal eukaryotes, which influence microbial food webs and nutrient cycling, signals a gap in research (Geisen et al., 2018; Tedersoo et al., 2014).

The evaluation of microbial communities as a function of biomass has been widely explored in numerous studies (Fig. 7)(Bargali, 2024; Xu et al., 2020). Assessing microbial communities through biomass provides a tangible and realistic measure of their contribution to ecological and biochemical processes (Smith & Paul, 2017; Bastida et al., 2021). However, biomass-based assessments alone often fall short in capturing the full extent of microbial diversity, particularly in terms of species richness, which is better determined through advanced sequencing technologies. Studies have shown that while biomass measurements provide quantitative data on microbial contributions, they cannot sufficiently uncover the functional and phylogenetic complexities of microbial ecosystems. Therefore, integrating biomass assessments with sequencing approaches is crucial for a comprehensive understanding of microbial dynamics (Fierer et al., 2020; Carter et al., 1999).

The majority of studies have focused on samples collected from urban forests, followed closely by urban parks (Fig. 8). This trend likely reflects the ecological significance of urban forests, as they often serve as remnants of natural forests, providing a habitat for native species and maintaining biodiversity within urban landscapes (Cocroft et al., n.d.; Alvey, 2006). Urban forests are particularly valuable for studying soil microbial communities due to their relatively undisturbed conditions, which offer a closer representation of natural ecosystems (Wu et al., 2024). Urban parks, on the other hand, represent a mix of native and exotic plant species, offering insights into how these species influence soil microbial communities and ecological interactions (Kourtev et al., 2003, Wang et al., 2022). In contrast, playgrounds, bare lands, and other less-studied urban environments have received minimal attention, possibly due to their perceived lack of biodiversity or ecological value (Horta et al., 2024). However, these underrepresented sites could provide unique perspectives on the adaptability and resilience of microbial communities in highly anthropogenic settings.

Since 2016, there has been a significant increase in research focusing on the soil microbiome and its interactions with vegetation (Fig. 4). This surge in publications highlights a growing awareness within the scientific community of the critical role that soil microbial communities play in ecosystem functioning and plant health (Newberger et al., 2023; Markalanda et al., 2022). The increase is likely driven by advancements in sequencing technologies, which have made it easier to study microbial diversity and functions in detail. Furthermore, the rising global challenges of soil degradation, climate change, and food security have underscored the importance of understanding soil microbial dynamics to develop sustainable solutions (Bertola et al., 2021). Researchers have started employing a combination of traditional methods, such as substrate-induced respiration, and modern approaches, like high-throughput sequencing, to gain a comprehensive understanding of soil microbial ecology (Beugnon et al., 2021). This multidisciplinary approach reflects a paradigm shift in soil research, moving beyond basic assessments to explore complex interactions within soil microbial communities and their impact on vegetation and ecosystem services (Liu et al., 2017).

Collectively, these highlight significant evidence gaps, such as the limited understanding of factors influencing soil microbiomes, insufficient focus on species diversity within vegetation, and an overemphasis on microbial biomass rather than utilizing sequenced data for more comprehensive analysis. There is a clear lack of studies being conducted globally on this topic, with only 62 relevant papers to date. While one can argue the relevance of this question, as mentioned above, with a growing urban population, more care needs to be taken regarding the type of vegetation present in urban spaces and the effect they ultimately have on soil microbes and, therefore, human health. Fig.1 shows the variety of factors that can affect soil microbiota, and several of these factors rely on urban management practices, including but not limited to the type of vegetation that is curated in such spaces. Urban green spaces frequently interact with humans and can directly influence their overall health and well-being. Overcoming methodological challenges and focusing on different aspects of soil health can promote better disease management and human health.

There is a general lack of research on urban space-based vegetation in most countries aside from North America, China, and Australia. Correlation studies between vegetation and microbial communities from other countries tend to focus mainly on agricultural spaces or along urban-rural gradients. Additionally, some of the initially shortlisted studies did not report on microbial indices or urban space vegetation. Furthermore, there is a significantly low number of studies, globally, that focus on vegetation diversity or species composition. This makes it difficult to draw effective conclusions about the effect of vegetation on soil if the vegetation parameters are not defined in detail. Several studies that focussed on microbial density only reported whether the microbial community belonged to bacteria or fungi. A deeper understanding of the kind of microbes that inhabit urban spaces can help us investigate this problem further in future studies.

## Limitations of the paper

We restricted our research to the English language, excluding non-English papers, which may have influenced our findings. However, we do not anticipate a significant bias in our evidence map, as only three papers in foreign languages were excluded. Additionally, some papers were excluded because they did not meet our inclusion criteria; for instance, they were not conducted in urban spaces, did not report on microbes, or did not address vegetation. While we acknowledge that these exclusions may introduce some bias, we have explicitly outlined our search criteria in the methods section to ensure transparency. Furthermore, we were unable to access the full-text versions of 10 papers, which were therefore excluded from the study. During the screening process, there was an agreement of 89.1% among reviewers on the inclusion and exclusion of papers. Disagreements were resolved through post-screening discussions to ensure consistency and accuracy.

## Conclusions

This systematic map is based on the comprehensive and systematic screening of available information on the impacts of above-ground vegetation on the soil microbiome community across diverse green spaces within urban areas. It identifies a wealth of information on the impact of vegetation communities that could conceivably affect the soil microbiome. As such it should be of value to a range of actors, including urban green space managers, researchers, and policymakers. As it is challenging for practitioners to read and synthesise the evidence on individual interventions and biodiversity outcomes, the map provides a key to finding concrete guidance from published research.

## Implications for Policy Management

Recognizing the critical role of soil microbiomes in ecosystems and human health, some policy recommendations have been proposed to integrate soil microbiome considerations into broader health and environmental strategies.

1. Inclusion in One Health Policies emphasizes the interconnectedness of human, animal, and environmental health. Achieving One Health objectives requires incorporating soil microbiomes into policy frameworks. An integrated science–policy–society approach can holistically address health challenges (Singh et al., 2023).
2. The development of Soil Health Indicators involves creating standardized metrics for soil health, including microbiome diversity and functionality, to guide land management policies and practices (Soil Health Institute, 2025). These benchmarks can assess the interventions’ impacts on soil and human health.
3. Public Awareness and Education efforts aim to foster community engagement in soil conservation through educational initiatives, empowering stakeholders to adopt practices supporting soil and environmental health (“The Soil Microbiome: A Game Changer for Food and Agriculture,” 2022). Implementing these policy changes requires collaboration among scientists, policymakers, and communities to effectively integrate soil microbiomes into strategies promoting health and sustainability.

## Implications for Research

The study of soil microbiomes holds transformative potential across multiple domains.

1. In agriculture, research drives the development of biofertilizers and biopesticides, offering sustainable alternatives to chemical inputs while enhancing crop productivity and soil health (FAO, 2021).
2. In biotechnology, understanding soil microbial diversity aids in discovering novel enzymes, antibiotics, and other bioactive compounds with applications in medicine, industry, and environmental remediation (Exploring Linkages Between Soil Health and Human Health, 2024).
3. In climate science, soil microbiome studies advance carbon sequestration strategies and greenhouse gas mitigation efforts, contributing to climate change solutions (Singh et al., 2023).
4. Furthermore, interdisciplinary research links soil microbiomes and human health, paving the way for microbiome-based therapies and nutritional innovations, such as microbiome-enriched agricultural products (Soil Health Institute, 2025).

Addressing current limitations, such as standardized methodologies and comprehensive databases, is essential for unlocking soil microbiomes’ potential in promoting sustainable development and human well-being.

The intricate relationship between above-ground vegetation and soil microbiomes underscores the importance of urban green spaces in sustaining ecosystem health and human well-being. This study emphasizes the need for a multidisciplinary approach that integrates research, policy, and public engagement to address critical gaps in our understanding and application of soil microbiome science. By leveraging advanced technologies, fostering global collaborations, and aligning conservation efforts with urban planning, we can unlock the full potential of soil microbiomes to combat pressing challenges such as biodiversity loss, climate change, and public health issues. Moving forward, a concerted effort to prioritize research and actionable strategies in this field will not only enhance ecological resilience but also ensure a sustainable and healthier future for urban communities worldwide.

## Supplementary Figure

**Supplementary Fig.1:**
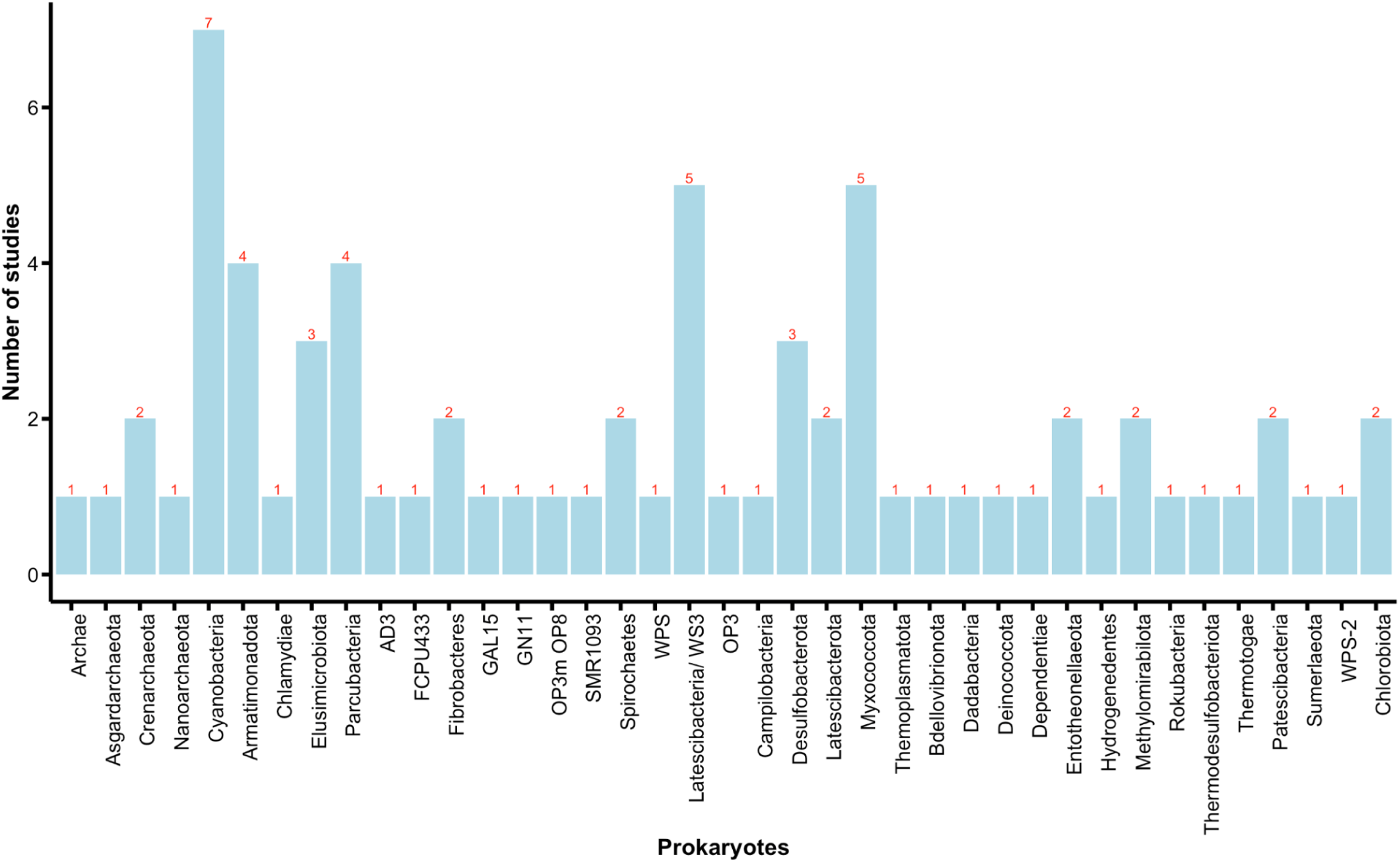
The bar chart shows the different families of prokaryotes observed in 10 or lesser studies.

**Supplementary Table 1:** The link to the table indicating the final list of studies used to make the evidence map. https://docs.google.com/spreadsheets/d/1z3HRK3KURNIS_B19fyHOR8xJ4hswNMHRdV0kIQGwcU/edit?usp=sharing

## Notes

### Competing Interest Statement

The authors have declared no competing interest.

